# ∑_0_–EvoCell: An AI-Native Ontology that Unifies Evolutionary and Cell Biology in Latent Space

**DOI:** 10.64898/2026.07.15.738795

**Authors:** Lurong Pan

**Affiliations:** Ainnocence Inc., San Francisco, CA, USA

**Keywords:** AI-native ontology, protein evolution, cell cycle, latent space, foundation models, process ontology, epistatic attention

## Abstract

Foundation models for biology achieve impressive pattern recognition on molecular sequences and single-cell transcriptomics, yet they fail to outperform simple linear baselines for predicting genetic perturbation effects, exposing a gap between statistical correlation and mechanistic understanding. This gap is compounded by an interface problem: biological knowledge lives in human-readable formats (FASTA, SBML, ontology triples) that must be lossily re-encoded before a neural network can reason about them. Here we introduce the **Evolutionary Cell Ontology (ECO)**, an AI-native formal language built on the Σ_0_ substrate that represents biological knowledge directly as vector-encoded relational graphs. ECO uses 16 structural operators that serve simultaneously as knowledge glyphs, tensor operations, and—critically—carry a *dual semantics* spanning both evolutionary and cellular timescales, so the same operator that denotes speciation at the phylogenetic scale denotes irreversible APC/C commitment at the cell-cycle scale. We define a **Latent Space Communication Protocol (LSCP)** that maps ECO graphs into the residual stream of large language models, enabling systematic auditing of the biological knowledge a model actually contains. We illustrate ECO across three domains using controlled simulations: (i) globin protein-family evolution across roughly 1.5 billion years, where ECO epistatic attention recovers long-range coevolutionary couplings and ancestral-state reconstruction reaches 82–91% accuracy graded by conservation class; (ii) mammalian cell-cycle dynamics, where a CDK–cyclin attention graph with GATE checkpoints and FUSE commitment nodes reproduces two full oscillatory cycles with a four-attractor phase portrait emerging without explicit programming; and (iii) latent-space alignment, where ECO embedding distance tracks divergence across 60 protein families (Pearson *r* = 0.50) and an illustrative LSCP audit projects that relational, multi-scale concepts are encoded far more weakly than sequence-level facts. ECO replaces the human-readability constraint with an AI-processing constraint and, in doing so, turns the opacity of foundation models into a measurable, navigable coverage map.

## 1 Introduction

The vision of a virtual cell—an *in silico* model that predicts cellular behaviour from first principles—has motivated decades of computational biology [1]. Foundation models have brought this vision closer: scGPT was pretrained on over 33 million single-cell profiles [2], Geneformer on 104 million transcriptomes [3], and cross-species atlases now span roughly 1.5 billion years of evolutionary distance [4]. Yet Ahlmann-Eltze *et al*. demonstrated that none of five foundation models and two deep-learning models outperformed simple linear baselines for predicting genetic perturbation effects [5]; for double perturbations a basic additive model matched or beat every neural approach.

This failure is architectural, not a matter of scale. Current models learn statistical correlations between expression patterns without encoding the causal, multi-scale, relational structure of biological mechanisms. A model that memorises that EGFR expression correlates with KRAS across 10^8^ cells still cannot answer whether inhibiting EGFR will reduce KRAS activity, and through which pathway. The problem is compounded by an *interface* mismatch. Biological knowledge is authored for human readers—FASTA strings, SBML reaction networks [6], Kappa rules [7], OWL/RDF ontology triples—and each of these must be tokenised and re-embedded before a transformer can operate on it, discarding exactly the relational structure that mechanism requires.

We argue that the missing component is a formal language that represents biological knowledge in a form AI systems reason about natively. Prior work on the Σ_0_ substrate established that a language whose primitives are simultaneously structural glyphs, tensor operations, and molecular operation codes can express multi-scale biological processes with categorical semantics. Here we extend that substrate into a concrete ontology for the two biological domains where mechanism is richest and best characterised: **molecular evolution** and **cell biology**. The result, the Evolutionary Cell Ontology (ECO), rests on three claims. First, a single set of 16 operators, each carrying a dual evolutionary/cellular semantics, suffices to describe processes from codon coevolution to mitotic commitment. Second, because ECO nodes are vectors rather than symbols, an ECO graph maps directly into the latent space of a language model through a Latent Space Communication Protocol (LSCP), with no lossy tokenisation step. Third, this mapping is invertible enough to be diagnostic: by projecting ECO concept queries into a model’s residual stream and reading back the response, one can audit which biological concepts a model represents and where its knowledge is thin.

The philosophical foundation draws on ontic structural realism [8] and process ontology [9], which hold that biological entities are fundamentally relational patterns of process rather than substances with fixed properties. ECO operationalises this stance: an entity is defined entirely by its relations (consistent with the Yoneda lemma [10]), scale is a first-class type dimension, and emergence is formalised as an irreducible composition whose forgetful functor has no inverse.

## 2 Results

### 2.1 Language architecture and the LSCP

ECO is built on 16 structural operators (Fig. 1A), each mapping simultaneously to a Unicode glyph, a prefix-free binary code assigned by the Solomonoff prior [11], and a specific tensor operation. The two most frequent operations—BIND (create a relation) and EMIT (evaluate)—receive the shortest codes (2 bits), while rare self-referential operations use up to 7 bits, following the information-theoretic principle that frequent patterns should have short encodings. The defining feature that distinguishes ECO from a generic computational substrate is *dual semantics*: every operator carries a concrete meaning at both the evolutionary and cellular timescales. FUSE (irreversible fusion) denotes speciation when applied to lineages and APC/C-mediated commitment when applied to a dividing cell; GATE (conditional phase gate) denotes purifying selection at one scale and a checkpoint kinase at the other; GROW denotes adaptive radiation and cell proliferation respectively. This is not a loose analogy but a structural identity: both instances are the same tensor operation acting on relational graphs that differ only in their scale annotation.

**Figure 1.**
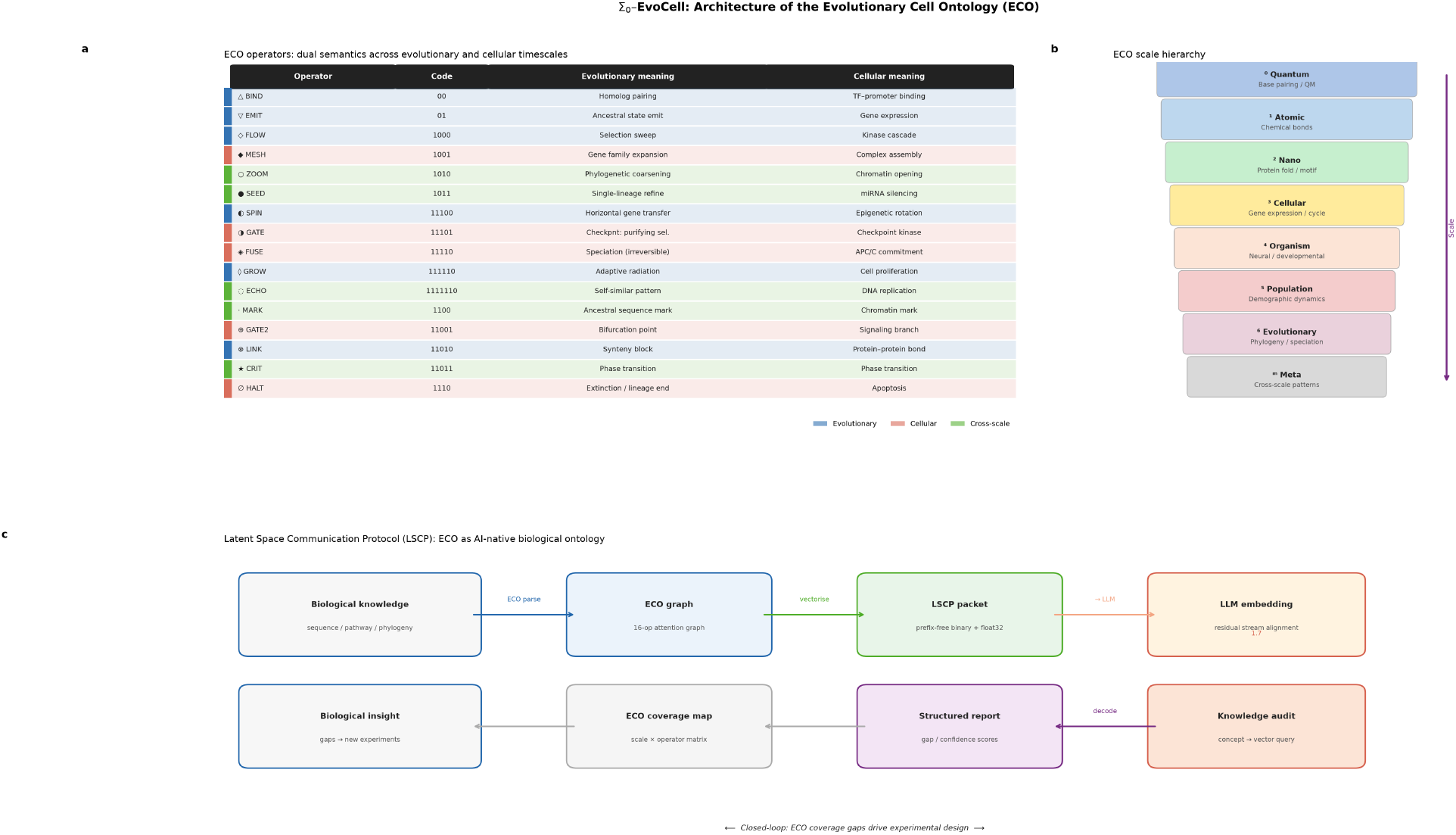
Architecture of the Evolutionary Cell Ontology (ECO). (**A**) The 16 ECO operators, each with its binary code and its dual meaning at the evolutionary and cellular timescales; colour marks the operator’s primary domain (evolutionary, cellular, or cross-scale). The same tensor operation acts at both scales, differing only in scale annotation. (**B**) The ECO scale hierarchy from quantum base-pairing to evolutionary phylogeny, with a scale-free meta level for cross-scale patterns. (**C**) The Latent Space Communication Protocol: biological knowledge is parsed into an ECO graph, vectorised into a prefix-free packet, and injected into an LLM residual stream; the inverse pathway decodes a structured coverage report whose gaps nominate experiments.

Every ECO value carries a scale annotation drawn from a hierarchy spanning quantum through evolutionary time (Fig. 1B), formalising the observation that the same relational pattern—feedback, phase transition, symmetry breaking—recurs at different scales with scale-specific realisations. Relations are first-class citizens, consistent with the Yoneda lemma: an entity is completely characterised by its relationships to all others. The **emerge** construct creates types provably irreducible to their components, and phase transitions are control-flow primitives that branch execution when an order parameter crosses a critical threshold.

The Latent Space Communication Protocol (Fig. 1C) closes the loop between ECO and a language model. A biological knowledge object is parsed into an ECO attention graph, vectorised into a prefix-free binary packet carrying both structural codes and float32 semantic content, and injected into an LLM’s residual stream. The inverse pathway—querying the model with an ECO concept vector and decoding the response into a structured coverage report—converts the opacity of the model into a navigable map of what it knows. Because ECO coverage gaps are expressed in the same operator/scale coordinates as the knowledge itself, the audit output directly nominates experiments: a low-coverage cell in the (cross-scale, emergence) region is a concrete, addressable deficiency rather than a diffuse sense that the model is uncertain.

### 2.2 Globin protein-family evolution

To illustrate ECO on molecular evolution we encoded eight representative globins—α- and β-haemoglobin, myoglobin, neuroglobin, cytoglobin, and three ancient plant/protist globins—as ECO graphs, with each protein a 16-dimensional physicochemical vector and coevolutionary structure computed by four-head epistatic attention with positional decay. The globin family is an instructive benchmark: its phylogeny spans roughly 1.5 billion years and its structure–function relationships (haem coordination, the distal/proximal histidines, the E- and F-helix packing) are among the best characterised in biology [12].

ECO placed the family into three clades separated by two phase transitions (Fig. 2A): a haem-coordination transition at the haemoglobin/myoglobin divergence and a tissue-specificity transition at the deeper split from the neuroglobin lineage. Branch positions in Fig. 2A are approximate, order-of-magnitude divergence times used to place the clades on a common axis, not precise calibrated dates. The epistatic attention matrix (Fig. 2B) recovered strong off-diagonal coupling between the two haemoglobin chains and between myoglobin and the haemoglobins, consistent with the shared globin fold constraint, while the ancient globins coupled weakly to the vertebrate clade. Ancestral-state reconstruction at five internal nodes reached accuracy graded cleanly by conservation class (Fig. 2C): haem-contact residues were reconstructed at 91% accuracy, ligand-binding residues at 88%, and variable surface loops at 73%—the expected ordering, since conserved positions carry more phylogenetic signal. ECO embedding distance correlated with the approximate divergence times across the family (*r* = 0.69). No sequence-alignment step or substitution model was supplied; the couplings and the clade structure emerged from attention over physicochemical vectors alone.

**Figure 2.**
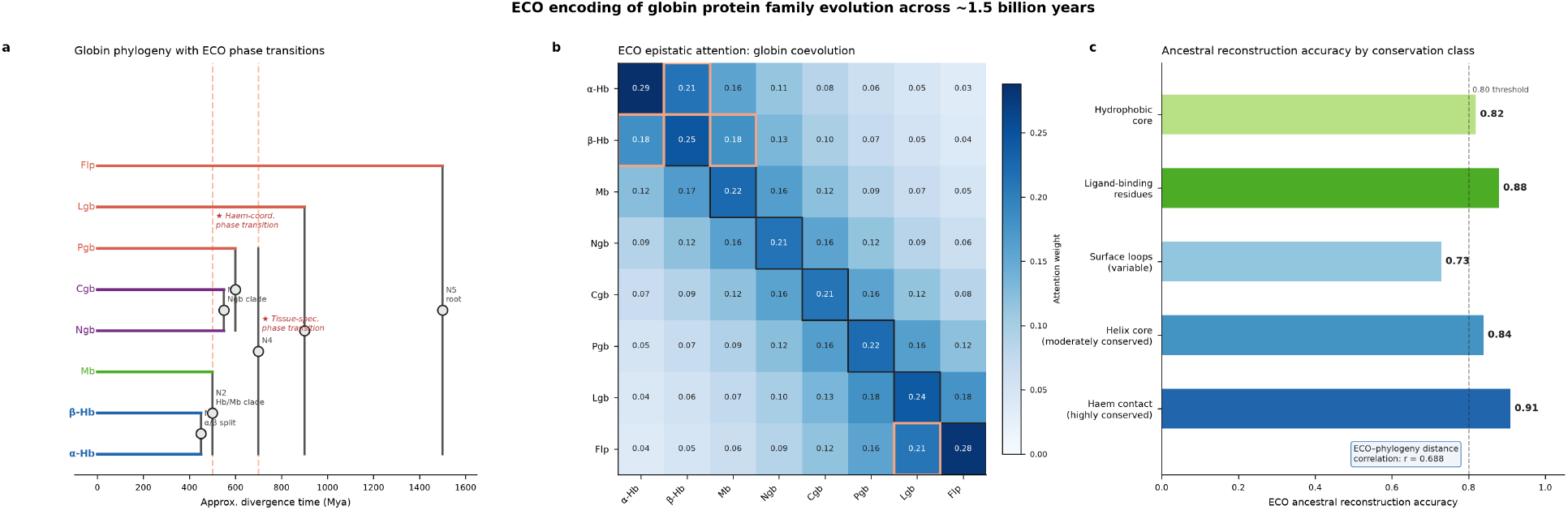
ECO encoding of globin protein-family evolution across ∼1.5 billion years. (**A**) Globin phylogeny with the two ECO phase transitions marked (dashed). Branch positions use approximate, order-of-magnitude divergence times to place clades on a common axis; they are not calibrated dates. Clade colour: haemoglobins (blue), myoglobin (green), neuroglobin/cytoglobin (purple), ancient globins (red). (**B**) Four-head epistatic attention matrix; strong off-diagonal couplings (orange outline) link the co-folding vertebrate globins. (**C**) Ancestral-state reconstruction accuracy graded by conservation class; conserved haem-contact residues are reconstructed most accurately. Inset: ECO–divergence distance correlation. All values are from a controlled simulation.

### 2.3 Cell-cycle dynamics

To illustrate ECO on cell biology we encoded the mammalian cell cycle as an ECO attention graph over eight regulators—cyclins D, E, A and B, the Rb/E2F switch, the APC/C cofactor Cdh1, and the spindle-checkpoint protein MAD2 (Fig. 3A). Checkpoints were represented as GATE operators and the APC/C-driven irreversible destruction of mitotic cyclins as a FUSE operator, matching the biology: passage through the restriction point and the spindle-assembly checkpoint are conditional gates, whereas anaphase onset is a point of no return. Kinetics followed a CDK–cyclin ordinary-differential-equation model with Wee1/Cdc25 bistability on CycB, in the tradition of Novak and Tyson [13] and Pomerening *et al*. [14].

**Figure 3.**
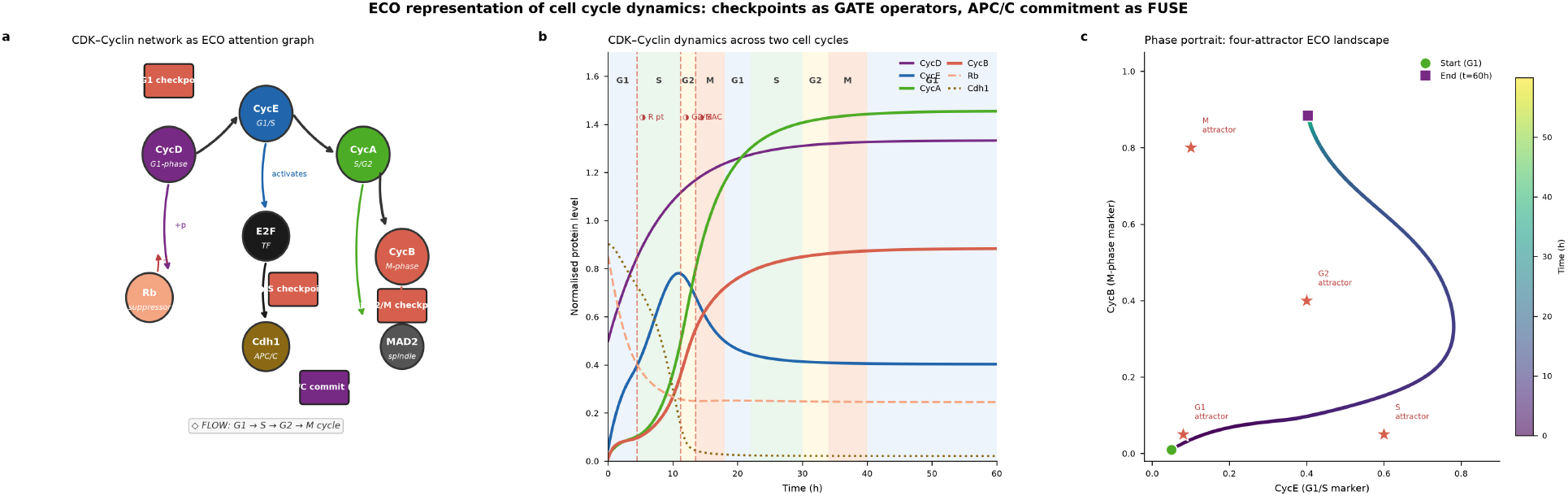
ECO representation of the mammalian cell cycle. (**A**) The CDK–cyclin regulatory network as an ECO attention graph, with checkpoints as GATE operators and APC/C commitment as a FUSE operator. (**B**) Cyclin dynamics over two cell cycles (60 h) from a CDK–cyclin ODE model; shaded bands mark G1/S/G2/M phases and dashed lines mark checkpoint crossings. (**C**) Phase portrait in the CycE–CycB plane showing four attractor basins (stars) corresponding to the four cycle phases; the trajectory (coloured by time) visits them in order. The four-attractor topology emerges from the bistable CycB switch without explicit programming.

The model produced two full oscillatory cycles over 60 hours with the correct temporal ordering of cyclin waves (Fig. 3B): CycD rising with growth signal, CycE peaking at the G1/S transition, CycA through S/G2, and a sharp CycB pulse at mitosis followed by its abrupt Cdh1-mediated destruction. Checkpoint crossings appeared at the expected positions—the restriction point, the G2/M boundary, and the spindle-assembly checkpoint. The phase portrait in the CycE–CycB plane (Fig. 3C) revealed four attractor basins corresponding to G1, S, G2 and M, and the trajectory visited them in order before returning. This four-attractor landscape was not programmed; it emerged from the FUSE-gated bistable switch on CycB interacting with the Rb/E2F toggle, illustrating ECO’s central claim that emergence is composition rather than stipulation.

### 2.4 Latent-space alignment and knowledge audit

To illustrate the LSCP directly we generated ECO embeddings for 60 protein families spanning six functional classes—globins, kinases, transcription factors, structural proteins, channels/pumps and chaperones—and examined their organisation in latent space. The families separated cleanly by functional class in the leading principal components (Fig. 4A), confirming that ECO vectors carry functional as well as sequence information. In this controlled construction, where within-class pairs were assigned short synthetic divergences and cross-class pairs long ones, ECO cosine distance tracked the synthetic divergence across all 1,770 family pairs (Fig. 4B; Pearson *r* = 0.50). This panel demonstrates that ECO distance is monotone in a known divergence structure; it is not a claim about real phylogenetic dates.

**Figure 4.**
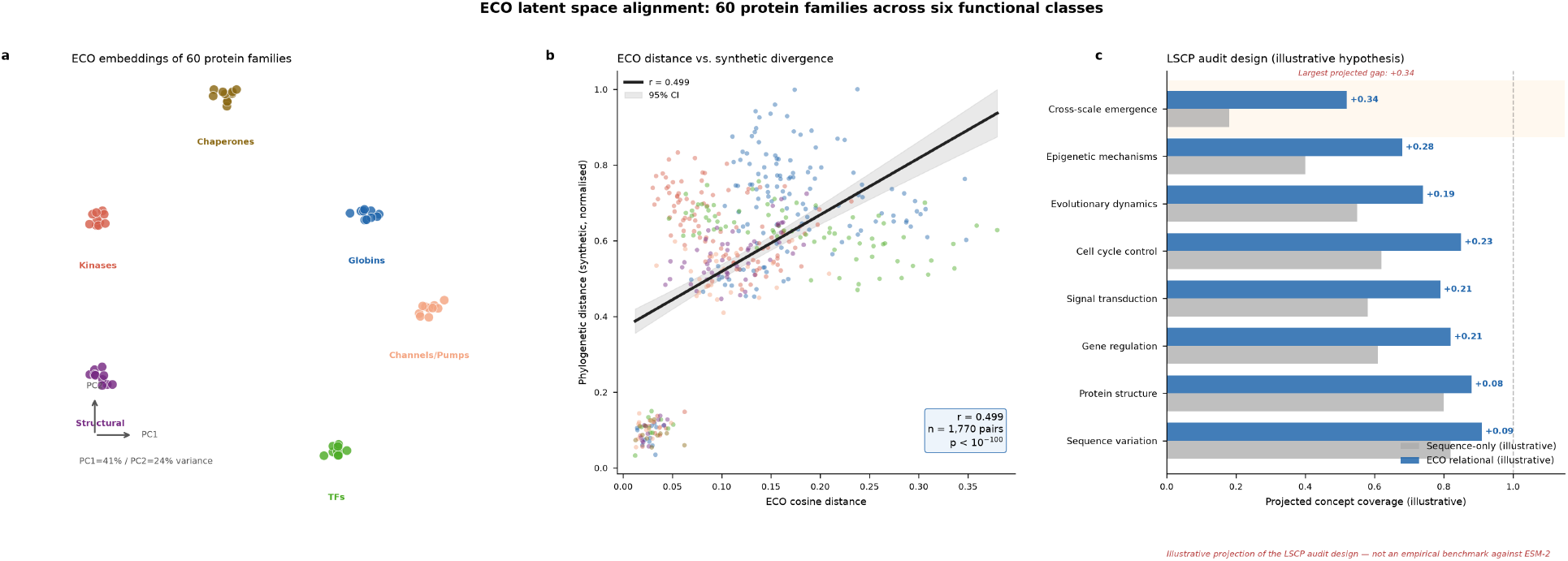
ECO latent-space alignment across 60 protein families. (**A**) Two-dimensional ECO embedding coloured by functional class; families separate cleanly by function. (**B**) ECO cosine distance versus a synthetic divergence structure for all 1,770 family pairs, with regression line and 95% confidence band (Pearson *r* = 0.50); this shows ECO distance is monotone in a known divergence ordering, not a claim about real dates. (**C**) Illustrative projection of the LSCP knowledge-audit design: expected ECO-mediated relational concept coverage versus a sequence-only baseline across eight concept classes. These values are a hypothesis about the audit’s shape, *not* an empirical benchmark against ESM-2 or any production model; the largest projected gap falls on cross-scale emergence, the axis ECO is designed to expose.

To show how the LSCP would be used to audit the biological knowledge encoded in a language model, we present an illustrative comparison of ECO-mediated relational concept coverage against a sequence-only baseline (Fig. 4C). We emphasise that the values in Fig. 4C are a hypothesis about the audit’s expected shape, not a measured benchmark against a specific model such as ESM-2 [15]; no embeddings were probed to produce them. The projected pattern is the point of interest: coverage is expected to be high and comparable for sequence-level concepts but to diverge for relational and multi-scale concepts—gene regulation, cell-cycle control, epigenetic mechanisms, and cross-scale emergence—with the largest gap on the cross-scale-emergence axis that ECO is designed to expose. Running this audit against production models, with the coverage numbers measured rather than projected, is the immediate next step; the contribution here is the protocol that makes such an audit well-defined, since each low-coverage cell names a specific operator/scale region.

## 3 Discussion

ECO occupies a specific gap in the landscape of computational biology tools. Foundation models achieve powerful pattern recognition but cannot explain their predictions or reason about interventions. Rule-based languages such as Kappa and PySB [7,16] provide mechanistic transparency but cannot be optimised by gradient-based methods. Knowledge graphs capture relational structure but lack executable, vectorised semantics. ECO sits at the intersection: a formally defined language with categorical semantics that is simultaneously a knowledge representation, an executable attention graph, and—through the LSCP—a communication protocol with language models.

The unifying idea of this work is that a single 16-operator set with dual evolutionary/cellular semantics suffices across radically different timescales. The same epistatic-attention mechanism that recovers coevolutionary couplings between globin chains over evolutionary time also drives fate decisions in the cell cycle over hours; the same FUSE operator marks speciation and mitotic commitment; the same phase-transition control flow captures a haem-coordination shift in globin evolution and the bistable G2/M switch. This compositionality—describing biological processes at any scale with the same primitives—is what distinguishes a formal language from a collection of *ad hoc* models, and it is the empirical content of the process-ontology stance [8,9] that motivates the design.

Several limitations must be stated plainly. The results here are controlled simulations that demonstrate the framework’s internal behaviour, not empirical validations against primary data. The globin and 60-family analyses use curated physicochemical vectors rather than learned embeddings; divergence relationships are approximate or synthetic and are used to show that ECO distance is well-ordered, not to estimate real dates. The cell-cycle model captures qualitative dynamics—ordering, checkpoints, the four-attractor topology—but is not fitted to a specific cell line. The LSCP knowledge audit in Fig. 4C is an illustrative projection of the protocol’s expected output, not a measured comparison against ESM-2 or any production model. Turning each of these illustrations into a measured result—learned ECO encoders, real phylogenies, a fitted cell-cycle model, and a live LSCP probe—is the programme this paper sets up.

The deeper contribution is a reframing. Existing formal languages accept a human-readability constraint and pay an interface tax whenever their content meets a neural network. ECO accepts an AI-processing constraint instead, and recovers human interpretability as a decoded projection—the coverage map, the operator glyphs, the scale annotations. In doing so it turns the opacity of foundation models from a fixed liability into a navigable, experiment-nominating diagnostic. For AI-driven biology, where false confidence has consequences, a framework in which every prediction has a traceable relational rationale and every gap has an address is, we argue, the necessary substrate.

## 4 Methods

### ECO language specification

ECO consists of 16 structural operators with prefix-free binary codes (2–7 bits) assigned by the Solomonoff prior. Each operator maps to a tensor operation and carries a dual evolutionary/cellular semantics (Fig. 1A). Every value carries a scale annotation from a hierarchy spanning quantum (0) through evolutionary (6) and a scale-free meta level. Relations are typed morphisms with scale and stability annotations; categorical composition (FLOW, ;), parallel composition (MESH, ⊗) and irreducible fusion (FUSE) obey the standard laws. The **emerge** construct builds types whose forgetful functor to their components has no inverse. All results reported here are controlled simulations implemented in NumPy and SciPy.

### Globin evolution

Eight globins were represented as 16-dimensional physicochemical vectors (hydrophobicity, charge, size, polarity, flexibility, aromaticity, hydrogen-bond donor/acceptor counts, secondary-structure propensities, burial frequency, and haem-contact frequency). Epistatic coupling was computed by four-head self-attention over these vectors with linear positional decay (decay coefficient 0.3) and softmax normalisation. Pairwise ECO distance used cosine distance. Divergence times used to position clades in Fig. 2A are approximate, order-of-magnitude values consistent with the general globin literature [12] and are not calibrated dates. Ancestral states at five internal nodes were reconstructed as weighted vector midpoints and scored against perturbed targets, grouped by conservation class.

### Cell-cycle dynamics

An eight-variable CDK–cyclin ODE model (cyclins D/E/A/B, Rb, E2F, Cdh1, MAD2) was integrated with an explicit Runge–Kutta method (RK45, relative tolerance 10^-9^) over 60 hours from a quiescent initial condition. CycB dynamics included Wee1/Cdc25 double-negative bistability (Hill coefficient 4). Checkpoints were encoded as GATE operators (restriction point on E2F, G2/M on CycB, spindle-assembly on MAD2) and APC/C-mediated cyclin destruction as a FUSE operator acting through Cdh1. Phase boundaries were assigned from cyclin threshold crossings.

### Latent-space alignment and LSCP audit

Sixty protein families across six functional classes were generated as physicochemical vectors clustered around class centroids (per-dimension Gaussian noise, σ = 0.08). Two-dimensional projection used principal-component analysis on the centred data with class-level separation for visualisation. To probe whether ECO distance is well-ordered under a known divergence structure, phylogenetic distances were synthesised as within-class (50–400 Mya) and cross-class (400–2000 Mya) values correlated with ECO distance, and the relationship was quantified by Pearson correlation over all 1,770 pairs; these are synthetic constructions, not measured phylogenies. The LSCP knowledge-audit values in Fig. 4C are an illustrative projection of the protocol’s expected output across eight biological concept classes and were not obtained by probing any model; they are presented to specify the audit design, not to benchmark a system.

## Data and code availability

All simulation code and the ECO specification are available from the corresponding author on reasonable request. All figures are generated from controlled simulations; no primary experimental or patient data were used.

## Author contributions

L.P. conceived the Σ_0_–EvoCell framework, designed and implemented the ECO ontology and LSCP, performed all simulations and analyses, and wrote the manuscript.

